# pH effect on strain-specific transcriptomes of the take-all fungus

**DOI:** 10.1101/2020.01.27.921023

**Authors:** Kévin Gazengel, Lionel Lebreton, Nicolas Lapalu, Joëlle Amselem, Anne-Yvonne Guillerm-Erckelboudt, Denis Tagu, Stéphanie Daval

## Abstract

The soilborne fungus *Gaeumannomyces graminis* var. *tritici* (*Ggt*) causes the take-all disease on wheat roots. Ambient pH has been shown to be critical in different steps of *Ggt* life cycle such as survival in bulk soil, saprophytic growth, and pathogenicity on plants. There are however intra-specific variations and we previously found two types of *Ggt* strains that grow preferentially either at acidic pH or at neutral/alkaline pH; gene expression involved in pH-signal transduction pathway and pathogenesis was differentially regulated in two strains representative of these types. To go deeper in the description of the genetic pathways and the understanding of this adaptative mechanism, transcriptome sequencing was achieved on two strains (PG6 and PG38) which displayed opposite growth profiles in two pH conditions (acidic and neutral). PG6, growing better at acidic pH, overexpressed in this condition genes related to energy production and protein deubiquitination. In contrast, PG38, which grew better at neutral pH, overexpressed in this condition genes involved in fatty acids metabolism. This strain also expressed stress resistance mechanisms at both pH, to assert a convenient growth under various ambient pH conditions. These differences in metabolic pathway expression between strains at different pH might buffer the effect of field or soil variation in wheat fields, and explain the success of the pathogen.

## Introduction

The filamentous fungus *Gaeumannomyces graminis* var. *tritici* (*Ggt*) is an ascomycete of large economic importance due to its devastating impact on cereal plants in temperate climates. The take-all disease caused by this fungus affects the roots of the host plants by blocking the conductive tissues and reducing water uptake. Serious infections under favorable conditions can result in decreased yields of up to 40%-60% [1].

*Ggt* populations are divided into two major genetically different groups (G1 and G2) which are known to coexist at the field scale in pluri-annual wheat monoculture experiments [2]. Ratios of G1 to G2 are different due to wheat crop history and disease level. G1 strains are more frequent in the first year of wheat monoculture, whereas G2 strains increase and reach a peak after three to five years corresponding to the maximum of take-all symptoms [3]. Furthermore, *in vitro* plant assays showed that G2 strains are slightly more aggressive than the G1 [4].

As most of soilborne pathogenic fungi, *Ggt* developed strategies to adapt to the ambient environmental factors all along its life cycle: survival in bulk soil, hyphal growth in soil during the saprophytic phase, and infection of host plants during pathogenic phase. This is particularly true concerning the pH factor. Soil pH is a factor influencing the take-all severity, whether modified by nitrogen supply [5] or by microbial communities interacting with *Ggt* and wheat [6], leading to a more acidic rhizosphere unfavorable to *Ggt* [7]. As a consequence, the pH-signaling pathway (Pal), characteristic of the fungal kingdom, has been shown to operate in *Ggt* [8]. This Pal pathway, first identified in *Aspergillus nidulans* [9], is composed of six proteins (palA, palB, palC, palF, palH, palI) which conduct pH signal to the transcription factor pacC [10]. Three forms of pacC exist: the inactive full-length pacC form predominates in acidic conditions whereas, in neutral-to-alkaline conditions, two proteolytic cleavages (the first one pH-dependent) enable pacC to be functional as a repressor of acid-expressed genes and an activator of neutral-to-alkaline-expressed genes [10].

Within *Ggt* species, evidence of intraspecific variability in pH sensitivity was demonstrated: some strains grew better at neutral pH and other at acidic pH [11]. More precisely, G1 and G2 strains are known to respond differently to the pH factor: whereas G1 and G2 strains have similar growth rate profile on acidic medium, G2 strains present a significantly better growth rate on neutral medium [11]. The mechanisms underlying the differential response of *Ggt* strains to the pH variations of the environment are not yet elucidated but extracellular pH has been shown to regulate *Ggt* gene expression involved in pathogenesis and saprophytic growth [8], in an original strain-specific way. Thus, the transcription factor pacC has been suspected to potentially play a role in pathogenesis through the regulation of expression of some pathogenesis-related genes that contained pacC binding sites (5’-GCCARG-3’). This is true for *Penicillium Expansum and Penicillium digitatum* in which pacC mutants are affected in their pathogenicity towards pear/apple and citrus fruits, respectively [12,13]. The ability of strains within the *Ggt* species to fine tune gene expression in response to the soil pH could affect growth rate leading to diverse (i) capacity of saprophytic growth (survival and development in bulk soil), (ii) capacity of surface roots’ colonization (first part of the pathogenic phase), and (iii) capacity of penetrating the roots (infection phase).

The aim of this study is to decipher *Ggt* gene expression regulation mechanisms involved in pH perception and to test the hypothesis of the link between pH perception and growth ability in the saprophytic growth and colonization of roots surface. Herein, two *Ggt* strains (PG6 and PG38), differing in their growth profile in function of the ambient pH and their group (G1 for PG6 and G2 for PG38), were selected and used for a RNA-Seq analysis under acidic or neutral pH conditions: pH 4.6 to mimick the value commonly found in soils and known to be unfavorable to *Ggt* [14], and pH 7.0 known as the optimal value for *Ggt* [5,15]. By this transcriptomics analysis, we showed that major metabolic and physiological changes associated with ambient pH and with pH-dependent growth of two strains occurred, and a focus was performed on the differentially expressed genes (DEGs) potentially involved in pH-dependent growth ability.

## Materials and Methods

### Fungal strains and culture conditions

*Ggt* strains used in this study (**Table 1**) were stored as potato dextrose agar (PDA) explants immersed in 10% glycerol at 4°C for long-term preservation. Prior to inoculation, each strain was cultured twice for 7 days at 20°C in the dark on autoclaved non-buffered (pH 5.6) Fåhraeus medium [16], the composition of which is described in the **S1 Table** [8]. Two Fåhraeus media buffered at pH 4.6 (A for acidic pH) or 7.0 (N for neutral pH), with different ratios of citrate / phosphate solutions (**S1 Table**), were then used for fungal mycelium growth measurement. pH was checked in all the media after autoclaving (115°C, 20 min). An Isopore 0.22 µm pore-size sterile membrane (Millipore, Molsheim, France) was laid on the agar surface to force mycelium to grow on top of the medium and to facilitate its sampling for further RNA extractions. Five mm diameter plugs of mycelium were removed from the edge of a colony grown twice the non-buffered medium. One plug per plate was laid on buffered Fåhraeus media (A or N), in the centre of Petri dishes, and the plates, covered by the polycarbonate filter, were incubated for 7 days at 20°C in the dark. Two orthogonal diameters of the colony were measured in each condition 7 days after inoculation. For each pH condition and strain, three plates were used, and three independent experiments were performed. Mycelium growth was compared using the Wilcoxon rank sum test.

**Table 1.** Origin of *Gaeumannomyces graminis* var. *tritici* isolates used in this study.

| Isolate | Original nomenclature | Molecular characterization (G1/G2) | Geographical origin / Isolation date | Source or reference | Host of origin |
| --- | --- | --- | --- | --- | --- |
| PG6 | 1125-6 | G1 | Germany / 1997 | Monsanto collection | Wheat |
| PG38 | 82/02 | G2 | France (Pacé) / 2002 | Lebreton et al., 2007 [3] | Wheat |

### RNA extraction from fungal mycelium grown on buffered media

After incubation at 20°C for 7 days, all the mycelium grown on polycarbonate membrane was collected. The mycelia from 3 plates were pooled and ground to powder with a pestle in liquid nitrogen-chilled mortars with Fontainebleau sand. Total RNA was extracted in 1 mL of Trizol (Invitrogen, Paisley, UK) and contaminating DNA was removed by using the RNase-free RQ1 DNase (Promega Corp., Madison, WI, USA) according to the manufacturer’s instructions. The RNA purity and quality were assessed with a Bioanalyser 2100 (Agilent Tech. Inc., La Jolla, CA, USA) and quantified with a Nanodrop (Agilent).

### Library construction, RNA-sequencing, and quality control

For each of the twelve samples (2 strains * 2 pH conditions * 3 biological replicates), sequencing libraries were generated using 2 µg of total RNA per sample with the TruSeq RNA sample preparation protocol from Illumina according to the manufacturer’s recommendations. Tags were added to each sample for identification. The sequencing reaction was performed using the Illumina HiSeq v3 chemistry on a HiSeq 2000 in 100 bp single read run according to the manufacturers recommendations (GATC Biotech, Konstanz, Germany). Sequence data quality control was evaluated using the FastQC program. Adapter sequences were removed using Flexbar, and Sickle was used to clean bases with substandard quality (PHRED 28) before removing reads under thirty bases length. Raw reads are available at the European Nucleotide Archive database system under the project accession number PRJEB34060.

### Read mapping to the reference genome and transcript counting

STAR v2.5.2a_modified [17] was used to align the reads to the published genome of *Ggt* strain (https://fungi.ensembl.org/Gaeumannomyces_graminis/Info/Index). This available reference sequenced genome is from the R3-111a-1 *Ggt* strain, different from the strains used in our study [15,18]. FeatureCounts v1.6.0 [19] was used to count the number of reads on each annotated *Ggt* gene, giving the raw counts. Genes with low count levels (under 1 count per million of mapped reads in three samples at least) were removed from the data.

### Differential expression analysis and gene clustering

DESeq2 R package [20] was used to product lists of DEGs between two experimental conditions. This program has its own normalization method “Relative Log Expression” (RLE) and needs to have raw counts as input. The p-values were adjusted to control multiple testing using the Benjamini and Hochberg’s method. Genes with an adjusted p-value < 0.05 were considered as significantly differentially expressed between conditions.

The clustering of gene expression profiles was performed using HTSCluster R package [21] by making groups of co-expressed genes on the DESeq2 normalized counts.

### Gene Ontology term (GO-term) enrichment

GO enrichment analysis of DEGs was achieved with TopGO R package [22] using weight01 algorithm, Fisher’s exact statistic test and a nodeSize parameter set to 5 (to remove enriched GO-term with less than five genes in the genome). GO-terms for each gene were first imported from BioMart database (Ensembl Fungi) via Blast2GO software (https://www.blast2go.com). For each GO category (Molecular Function, Cellular Component and Biological Process), Top 10 Enriched GO-terms (p-value < 0.05), enrichment ratios (> 5), and number of genes under each enriched GO-term were represented using ggplot2 R package.

### Quantitative real-time PCR (qRT-PCR) validation

The expression levels of 8 selected DEGs were determined by qRT-PCR to confirm the results of RNA-Seq analysis. Total RNA from *Ggt* mycelium was reverse transcribed with a set of two external RNA quality controls as previously described [8,23]. Briefly, for all the strains and the pH conditions, 750 ng of total RNA from fungi were mixed with known quantities of the two external controls. Reverse transcription was carried out in 30 µL containing 375 ng of random primers, 1 X ImPromII reaction buffer, 3 mM MgCl_2_, 125 µM of each dNTP, 30 U of RNasin Ribonuclease Inhibitor and 1.5 µL of ImProm-II™ (Promega). The following parameters were applied: 5 min at 25°C, 1 h at 42°C and 15 min at 70°C. Reactions without RNA or without reverse transcriptase were performed as negative controls. The oligonucleotides designed with the Primer 3 software are described in **S2 Table**. Quantitative PCR reactions (20 µL) containing 1 µl of cDNA, 0.4 µM of each primer and 1 X SybrGreen I Master (Roche) were performed on the LightCycler® 480 Real-Time PCR System (Roche). The quantitative PCR profile consisted of an initial denaturation at 95°C for 5 min, followed by 45 cycles of 95°C for 15 sec and hybridization-elongation temperature for 40 sec (**S2 Table**). A dissociation stage was applied at the end of the PCR to assess that each amplicon generated was specific. Moreover, each specific amplicon was sequenced (Genoscreen, Lille, France) to confirm it corresponds the expected sequence. The expression levels of transcripts were normalized using external RNA controls [23,24] from three independent biological replicates, each with three technical PCR replicates. Data were analyzed using the ANOVA procedure of the R statistical analysis software.

## Results and discussion

### Effect of pH on growth

The two *Ggt* strains (PG6 and PG38) were grown on media buffered at pH 4.6 (A) or 7.0 (N) in the dark. After 7 days, two orthogonal diameters of the colony were measured in each condition (**Fig 1**). Both strains grow at both pH levels but displayed two different growth profiles. PG6 showed a significantly higher mycelium growth at the acidic pH compared to the neutral condition. On the contrary, PG38 grew significantly better in a neutral medium than in an acidic environment. These growth rate profiles are representative of the interspecific variability of *Ggt* populations [8,11]. The growth profiles were different between strains according to the pH making relevant the study of the molecular basis regulating these two distinct phenotypes.

**Fig 1.**
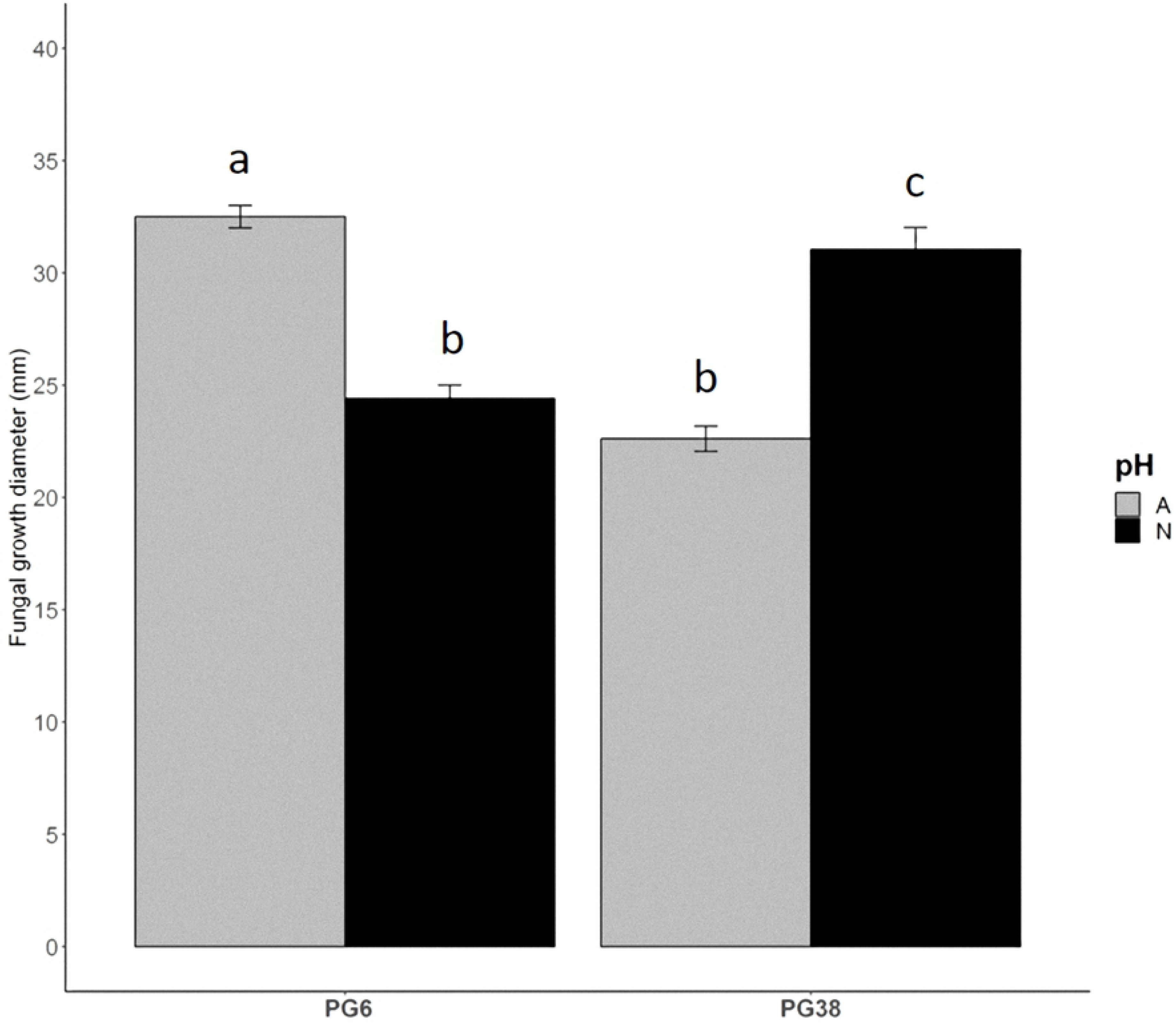
Mycelium growth of *Ggt* strains as a function of medium pH. PG6 and PG38 *Ggt* strains were grown at 20°C in the dark on media buffered at pH 4.6 (A) or 7.0 (N). The *Ggt* plugs were laid on medium covered with a polycarbonate filter allowing the mycelium sampling. The colony diameters were measured after 7 days of incubation. Each value is the mean of three biological replicates and three technical replicates. Error bars represented standard errors of the means. Conditions with different letters were statistically different according to the Wilcoxon rank sum test (P<0.05). Strains were depicted on the x-axis and fungal growth diameter in mm was on the y-axis.

### Overview, mapping and counting of the RNA-Seq data

RNA-Seq expression profiling was performed on the two strains (PG6 and PG38) growing on two different pH (4.6 or 7.0). The three biological replicates of each condition were sequenced in one pool of 12 tagged libraries on an Illumina HiSeq 2000 in 100 bases single-read mode. From 11,956,072 to 20,507,372 raw reads were obtained per sample (**Table 2**). After removing adapters and low quality bases (< Q28), 91.0 to 92.6% of the raw reads were kept with a minimum of length set to 30 bases. Thus, a total of 10,957,477 to 18,783,248 cleaned reads per sample were generated from the different RNA libraries. After read trimming, 78.2 to 86.1% of the initial raw data were uniquely mapped to the *Ggt* genome. Reads with multiple location (0.2 to 1.0 %) and too short alignements (6.0 to 13.3%) were removed. At the end, 70.2 to 78.5% of the raw reads were kept. We remomved the raw counts with less than one count per million of reads in at least three samples. We kept 11,041 genes among the 14,744 contained in the *Ggt* genome. Thus, the accuracy and quality of the sequencing data were sufficient for further analysis

**Table 2.** Overview of number of raw, cleaned, mapped, and counted reads from the different samples.

| Sample name | Library for each replicate | Number of total raw reads | Reads after cleaning and trimming (I30, q28) |  |  |  |  |
| --- | --- | --- | --- | --- | --- | --- | --- |
|  |  |  | Number of cleaned reads | Number of uniquely mapped reads <sup>1</sup> | % | Number of counted reads <sup>2</sup> | % |
| PG6 / A | 1 | 15,659,005 | 14,506,452 | 12,640,293 | 80.7 | 11,430,087 | 73.0 |
|  | 2 | 12,994,429 | 11,874,270 | 10,225,348 | 78.7 | 9,126,369 | 70.2 |
|  | 3 | 15,355,211 | 14,211,595 | 12,249,379 | 79.8 | 10,991,518 | 71.6 |
| PG6 / N | 1 | 19,466,804 | 17,734,183 | 15,223,234 | 78.2 | 13,666,878 | 70.2 |
|  | 2 | 16,798,714 | 15,407,637 | 13,261,468 | 78.9 | 12,039,586 | 71.7 |
|  | 3 | 17,529,036 | 16,035,463 | 13,869,536 | 79.1 | 12,636,834 | 72.1 |
| PG38 / A | 1 | 17,641,336 | 16,047,490 | 14,867,129 | 84.3 | 13,654,777 | 77.4 |
|  | 2 | 17,583,033 | 16,111,331 | 15,082,270 | 85.8 | 13,762,219 | 78.3 |
|  | 3 | 14,107,598 | 12,956,289 | 12,150,819 | 86.1 | 11,071,449 | 78.5 |
| PG38 / N | 1 | 20,507,372 | 18,783,248 | 17,241,029 | 84.1 | 15,636,718 | 76.2 |
|  | 2 | 17,616,201 | 16,206,580 | 15,024,729 | 85.3 | 13,707,008 | 77.8 |
|  | 3 | 11,956,072 | 10,957,477 | 9,938,207 | 83.1 | 9,012,954 | 75.4 |

The percentages are given from initial raw reads.
<sup>1</sup> Total number of reads mapped to uniquely locations in the *Ggt* genome with STAR
<sup>2</sup> Total number of reads counted with FeatureCounts
A: acidic pH and N: neutral pH

### Validation and clustering of the RNA-Seq data

We looked for differential gene expression between the different samples and normalized DESeq2 counts of all samples were plotted on a PCA to estimate the variability of the experiments and the biological conditions (**Fig 2**). 86% of the variance was represented on the plot. We confirmed that the three replicates of each experimental condition were largely clustered together, which validate our RNA-Seq experiment. The x-axis (57%) separated clearly the two strains whereas the y-axis (29%) was representative of the pH effect.

**Fig 2.**
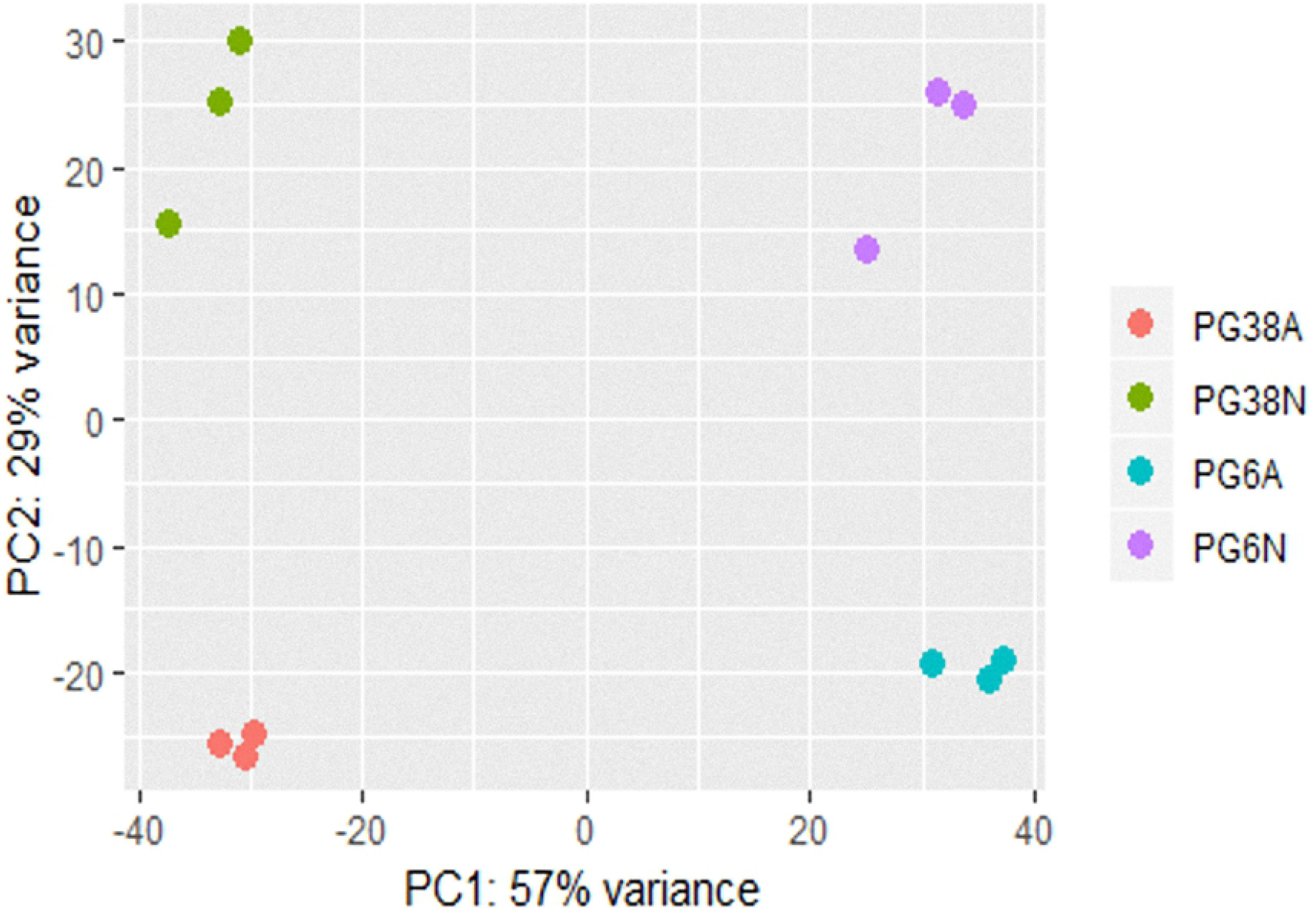
Estimation of biological variations by a Principal Component Analysis of the 12 transcript profiles. X-axis represented the variance on the first axis of the PCA, and the second was picted on the y-axis. A: acidic pH and N: neutral pH

As biological replicates were homogeneous, gene expression level of the four experimental conditions was calculated based on the means of the three replicates of each biological condition (**Fig. 3**). Expression profiles were clearly different between conditions, identifying groups of up-regulated (red) and down-regulated (green) genes. Gene expression was affected by the pH and/or by the strain. A clear separation was seen first between the strains, and secondly between pH inside each strain.

**Fig 3.**
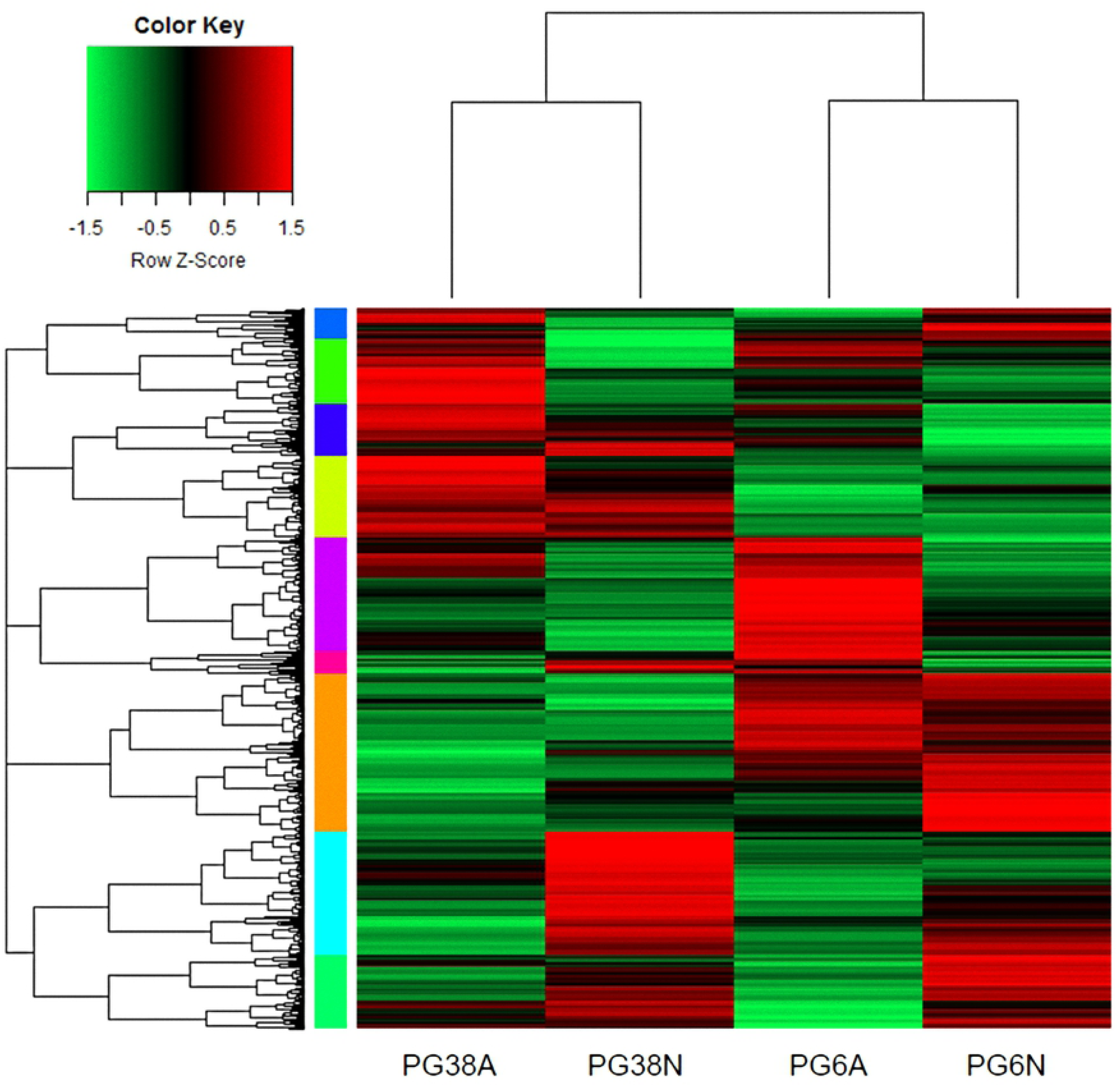
Overview of gene expression. Gene expression levels were displayed from green (downregulated) to red (upregulated). Colored bars on the left of the heatmap mark distinct major branches in the clustering tree grouping genes with similar expression pattern. Each row corresponded to a single gene and each column to the mean of the three biological repetitions of one experimental condition. The heatmap was generated with scripts based on heatmap.2 function as available in the “gplots” R package. A: acidic pH and N: neutral pH

In order to better characterize the different expression profiles of the expressed genes, we performed a co-expression clustering analysis based on Poisson Mixture models [21]. The analysis described 6 different profiles (**Fig. 4**). One profile (cluster 1) showed genes with strong expression in PG6, especially at acidic pH (1,167 genes) and another profile (cluster 2) showed genes with high expression in PG38 at neutral pH (445 genes). Both clusters 1 and 2 were representative of the higher growth rates of both strains according to pH. On the contrary, clusters 3 (1,168 genes) and 4 (503 genes) depicted the genes expressed in PG38 and PG6, respectively, in the pH where their growth were the lowest. In addition, clusters 2, 4, and 6 depicted genes more expressed at neutral pH compared to acidic pH: 445 genes highly expressed in PG38 compared to PG6 (cluster 2), 503 genes, including pacC, highly expressed in PG6 compared to PG38 (cluster 4), and 3,034 genes with similar expression in both strains (cluster 6). As the transcription factor pacC is known to repress acid-expressed genes and to induce neutral-to-alkaline expressed genes [10], the presence of pacC in the cluster 4 confirmed the activation of pH-signaling pathway at neutral pH in our study. The clusters 3 and 5 were more homogeneous between samples whereas cluster 3 showed genes globally more expressed in PG38 and cluster 5 in PG6 (4,724 genes). Interestingly, the clusters 3 and 5 contained the expression of the different pal genes confirming the expression of this pH-signaling pathway in *Ggt* [8].

**Fig 4.**
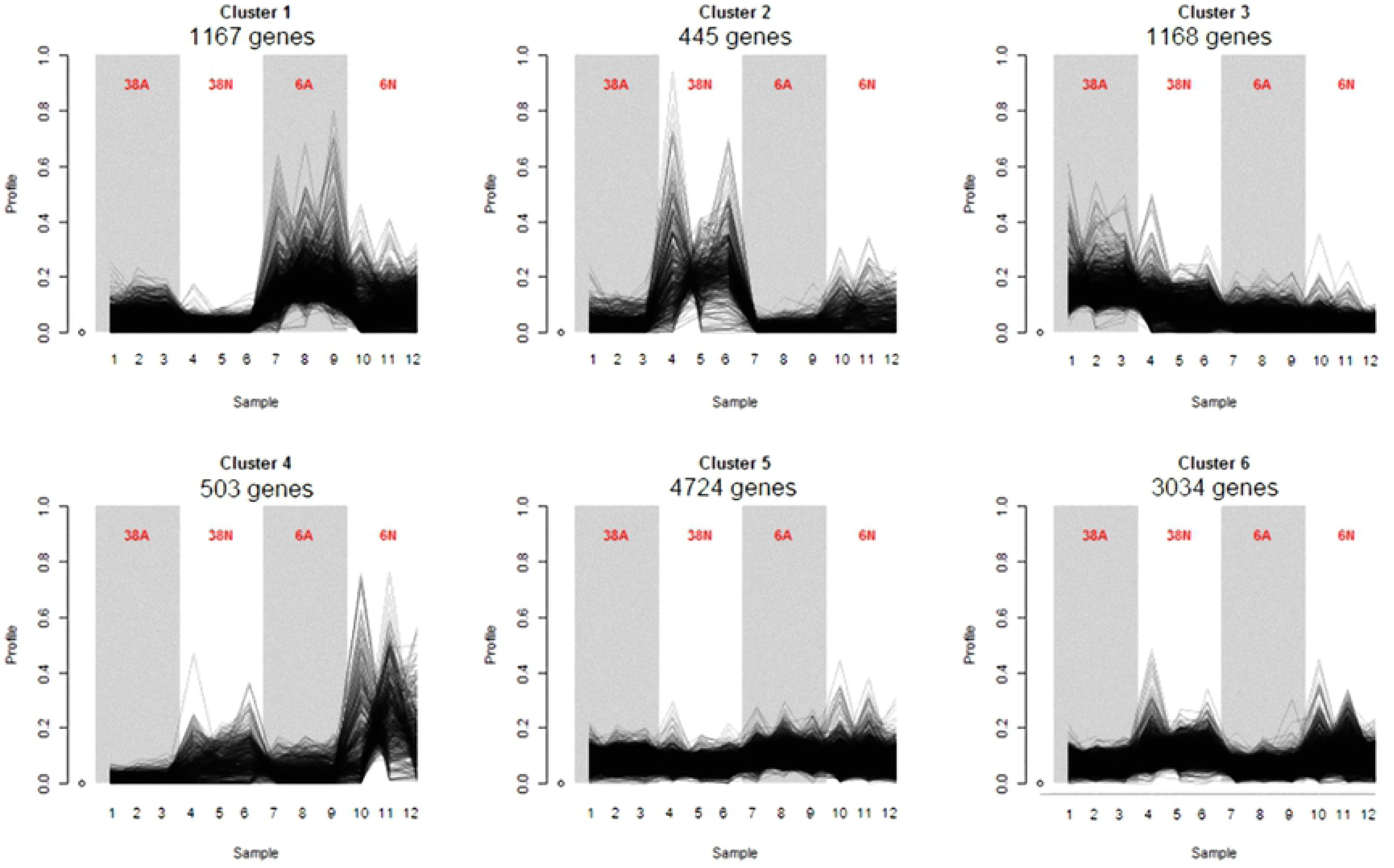
Co-expression clusters. HTSCluster was used to cluster expression data with Poisson Mixture Models. The number of genes assigned to a particular cluster was indicated below the cluster name. Grey and white backgrounds were for acidic and neutral pH, respectively. Each of the 12 samples was depicted on the x-axis and each biological condition (three replicates) was on the top of the curves. Y-axis represented, for each gene, the number of counts divided by the total number of counts in all the samples (frequency). A: acidic pH and N: neutral pH

Finally, the clusters summarized well the different biological growth profiles that took place according to the strains and the pH, as described in the **Fig 1**, making relevant the study of transcriptional profiles in relation to growth phenotypes.

### Overall comparison of differentially expressed genes (DEGs)

After DESeq2 normalization, four main comparisons biologically relevant were performed: 3,269 genes were differentially expressed between PG6A and PG6N conditions, 3,874 between PG6A and PG38A, 3,171 between PG38A and PG38N, and 3,651 between PG6N and PG38N (**Set Size on Fig. 5**). Each of these DEGs lists contained a similar number of genes and represented from 21.5 to 26.3% of the total number of genes of the *Ggt* genome. Among these lists, 605 genes were specific (not found in other comparisons) of the PG6A/PG6N comparison (**blue arrows on Fig. 5**), 655 of the PG6A/PG38A contrast (**red arrows on Fig. 5**), 544 of the PG38A/PG38N contrast (**green arrows on Fig. 5**), and 541 of the PG6N/PG38N contrast (**orange arrows on Fig. 5**). Among these four last lists, the percentages of up-regulated and down-regulated were similar and the total number of specific DEGs represented from 3.7 to 4.4% of the total number of genes included in the *Ggt* genome.

**Fig 5.**
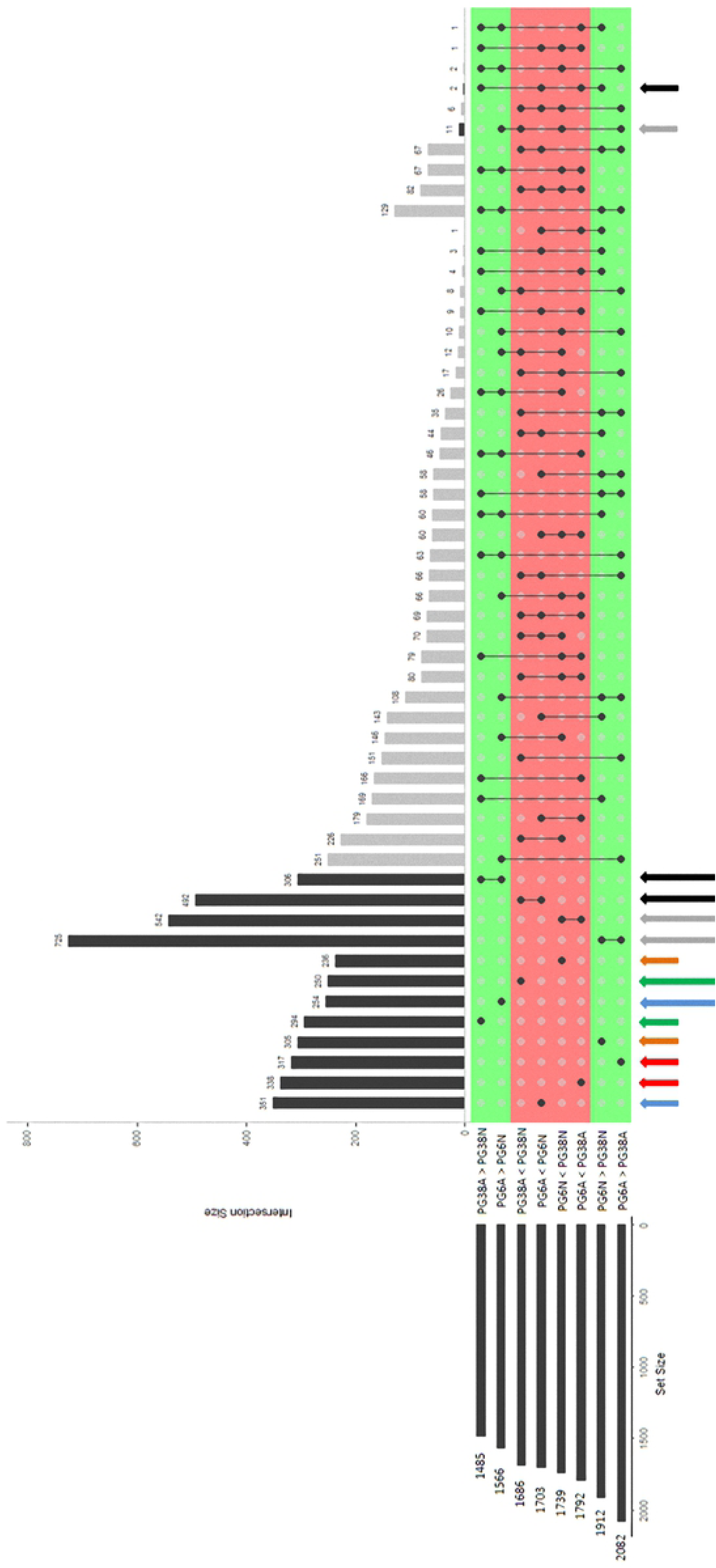
Plot of intersections between sets of genes differentially expressed. The intersected lists of differentially expressed genes and their sizes (number of DEGs) were presented in the horizontal bars on the left. The connected lines among the lists represented their intersections. The vertical bars and associated numbers corresponded to specific overlap of DE gene sets. The legends of colored arrows were explained in the text. The significance of gene expression changes was inferred based on an adjusted p-value < 0.05. <: number of DEGs underexpressed in the first condition compared to the second. >: number of DEGs overexpressed in the first condition compared to the second. A: acidic pH and N: neutral pH

The UpSetR package was used to identify 798 genes specifically regulated between acidic and neutral pH whatever the strain (**black long arrows on Fig. 5 and Table 3**), and 1,267 genes regulated between the two strains independently of the pH (**grey long arrows on Fig. 5 and Table 3**).

**Table 3.** Number of DEGs according to strain (A) and pH (B).

| A | Comparison | Up / Down | Number of DEGs |
| --- | --- | --- | --- |
|  | PG6 compared to PG38<br>(1,267 genes) | PG6 < PG38 | 542 |
|  |  | PG38 < PG6 | 725 |
| B | Comparison | Up / Down | Number of DEGs |
|  | Acidic pH compared to neutral<br>(798 genes) | A < N | 492 |
|  |  | N < A | 306 |

<: number of DEGs underexpressed in first condition compared to the second.
A: acidic pH and N: neutral pH

As both studied factors (strain and pH) had an effect on *Ggt* transcriptome, the study focused on transcriptomics differences between strains whatever the pH (strain effect) on one hand, and on transcriptomics differences between pH whatever the strain (strain effect) on the other hand. Finally, to highlight mechanisms potentially involved in the specific ability of each strain to grow under favorite pH, the overexpressed genes for each strain in the pH condition they grew better were more specifically analyzed.

### Strain effect on the *Ggt* transcriptome whatever the pH

To better understand how the transcriptome differed between the strains whatever the pH, a GO-term enrichment analysis was achieved with TopGO. Among the 1,267 DEGs linked to the strain effect (**grey long arrows on Fig. 5 and Table 3**), 542 and 725 were overexpressed in PG38 and PG6, respectively (**Table 3**). The **Fig. 6A and 6B** showed that 51 genes were involved in an enrichment of 27 significant GO-terms. Among these 27 enriched GO-terms related to the strain effect, 16 GO-terms were enriched in PG38 (**Fig. 6A**) and 11 in PG6 (**Fig. 6B**).

**Fig 6.**
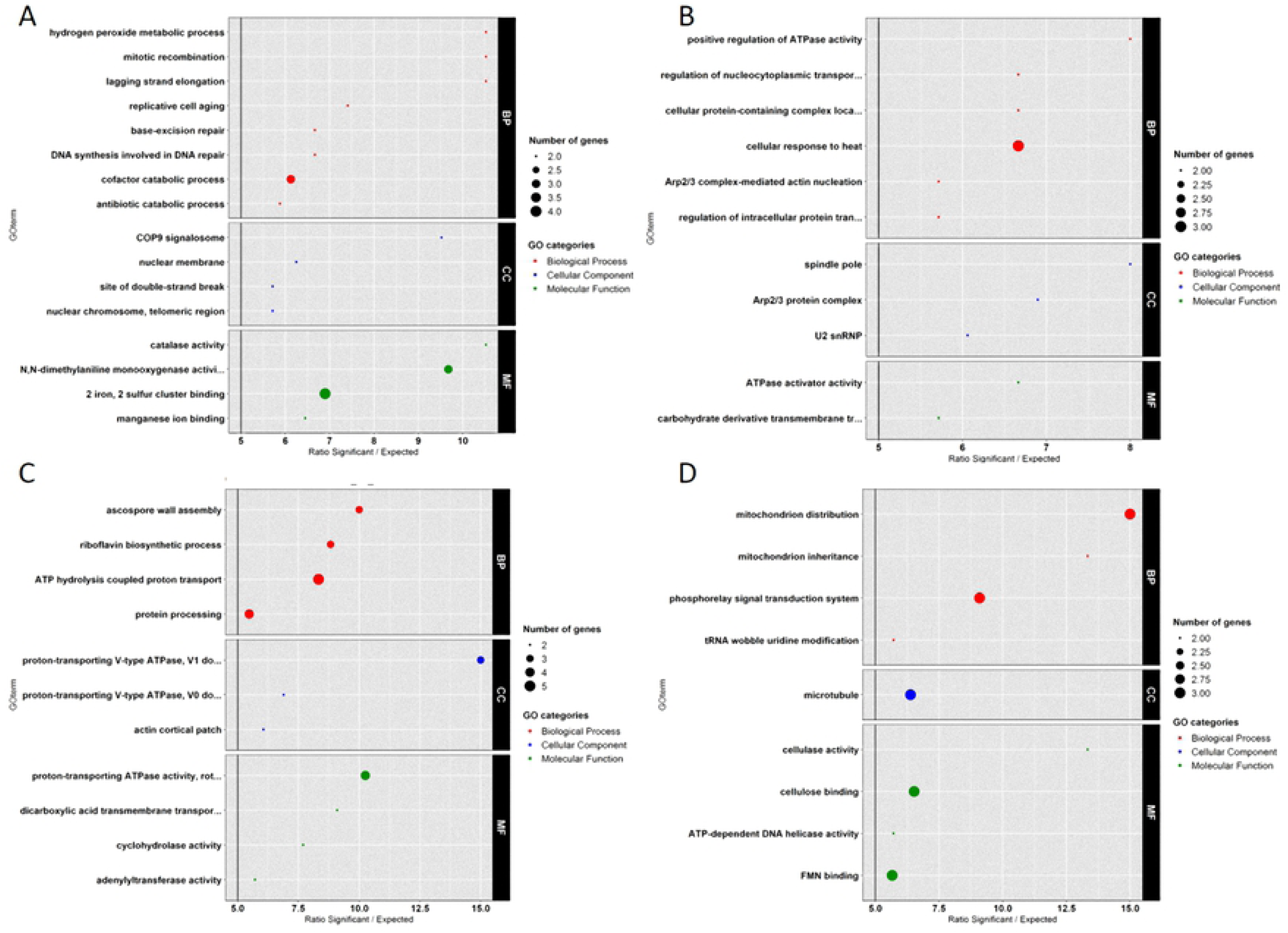
GO enrichment on DEGs lists according to strain or pH effects. Enrichment factor (> 5) was represented on the x-axis and enriched GO-terms were described on the y-axis. (A) Genes linked to strain effect: under-expressed in PG6 compared to PG38 (B) Genes linked to strain effect: over-expressed in PG6 compared to PG38 (C) Genes linked to pH effect: under-expressed in acidic medium compared to neutral (D) Genes linked to pH effect: over-expressed in acidic medium compared to neutral BP = Biological Process, CC = Cellular Component and MF = Molecular Function

The GO-terms enriched in PG38 were involved in DNA repair and DNA replication mechanisms (‘lagging strand elongation’, ‘replicative cell aging’, ‘base excision repair’, ‘DNA synthesis involved in DNA repair’, and ‘mitotic recombination’). The ‘base excision repair mechanism’ has been especially shown to be involved in host defense in the human pathogenic fungus *Paracoccidioides brasiliensis* [25]. The ‘2 iron, 2 sulfur cluster binding’ term was essential to cell viability in *Saccharomyces Cerevisiae* via its implication in ribosome assembly, DNA damage repair, and DNA replication [26]. In addition, the ‘antibiotic catabolic process’ enriched GO-term in PG38 could take part of a global stress-signaling pathway and could, for example, explain a potential ability of this strain to resist to antibiotics synthetized by antagonists microogranisms in soils [6]. So whatever the pH, the PG38 seemed to apply DNA repair and stress resistance mechanisms which could be essential in survival during wheat intercrop and growth of *Ggt* in soils and on roots surface.

On the other side, the enriched GO-terms in PG6 were associated to mitochondrial inheritance, energy transfer, and energy production. For example ‘Arp2/3 complex’ was essential to mitochondrial material transport to daughter cells during mitosis [27,28] participating in nucleation of actin filaments. As this transport was also supported by microtubules, the enrichment of ‘spindle pole’ GO-term in PG6 could be important for mitochondrial transport by its action on microtubule organization. Finally, ‘ATPase activity’ enriched in PG6, suggests an important role for energy transfer. In PG6, a large part of its metabolism seemed to be related to energy mechanisms potentially important for its growth whatever the pH.

### pH effect on the *Ggt* transcriptome whatever the strain

In the same way, TopGO analysis enabled to identify GO-term enrichments in the genes overexpressed at a given pH compared to the other pH, whatever the strain. Among the 798 genes previously described as linked to the pH effect (**black long arrows on Fig. 5 and Table 3**), 492 and 306 were respectively overexpressed in neutral and acidic environment. **Fig. 6C and 6D** showed that 53 genes of these DEGs were involved in the enrichment of 20 significant GO-terms. Among these 20 enriched GO-terms linked to the pH effect, 11 GO-terms were enriched in neutral conditions (**Fig. 6C**) whatever the strain and 9 in acidic conditions (**Fig. 6D**).

Several enriched GO-terms in neutral conditions were linked to ‘V-ATPases’ and ‘proton transport’. These pathways have been demonstrated as involved in organelles acidification in cells in response to extracellular pH, with an overexpression at pH 7.0 in yeast cells [29] as for *Ggt* in this study. Under these enriched GO-terms, we found 5 genes (GGTG_01849, GGTG_04071, GGTG_06975, GGTG_04610, and GGTG_10757) expressed at the same level between conditions N and A in the two strains and displaying similar fold change (about 1.5) between neutral and acidic conditions. So both strains of our study seemed to be able, at the same level, to maintain pH-gradients in their cells and organelles when they grew on neutral medium. Enrichment of ‘ascospore wall assembly’ at neutral pH was also seen, suggesting an ability of both strains at this pH to form ascospores. As this ability is related to the sexual stage of *Ggt*, which is usually used to disperse at long distance or to resist to stresses, it could have an importance in dynamics of the take-all disease during wheat monocultures, in function of the ambient pH.

On the other side, the main enriched function at acidic pH was linked to ‘mitochondria transport and inheritance’ (GO-terms ‘mitochondrion distribution’, ‘mitochondrion inheritance’, and ‘microtubule’). As the mitochondria activity could affect various mechanisms such ATP production, cellular differentiation, or cell death [27,28], the role of these enriched genes needs to be elucidated.

### Strain and pH effects on *Ggt* transcriptomes according to the growth profiles

To understand more precisely the mechanisms potentially involved in the specific ability of each strain to grow on its favorite pH, we focused on genes overexpressed in each strain in the pH condition they grew better.

Thanks to a TopGO identification of enriched GO-terms among the 250 genes overexpressed in PG38 at neutral pH compared to acidic (**Fig. 7 A** and **green long arrow on Fig. 5**), 41 contributed to 15 GO-terms enrichment. A main part of the enriched GO-terms (9) was involved in fatty acid metabolism : ‘fatty acid transport’, ‘fatty acid catabolic process’, ‘monocarboxylic acid catabolic process’, ‘acyl-CoA metabolic process’, ‘peroxisomal part’, ‘integral component of peroxisomal membrane’, ‘thiolester hydrolase activity’, ‘CoA hydrolase activity’, and ‘acyl-CoA hydrolase activity’. As fatty acids are the main source of carbon and energy in fungi [30], this pathway could play a role in the ability of PG38 to better grow at neutral pH. In addition, this strategy seemed to be specific to the growth of PG38 because fatty acid pathways were not enriched in the acidic condition in which PG6 showed better growth (**Fig. 7 B**).

**Fig 7.**
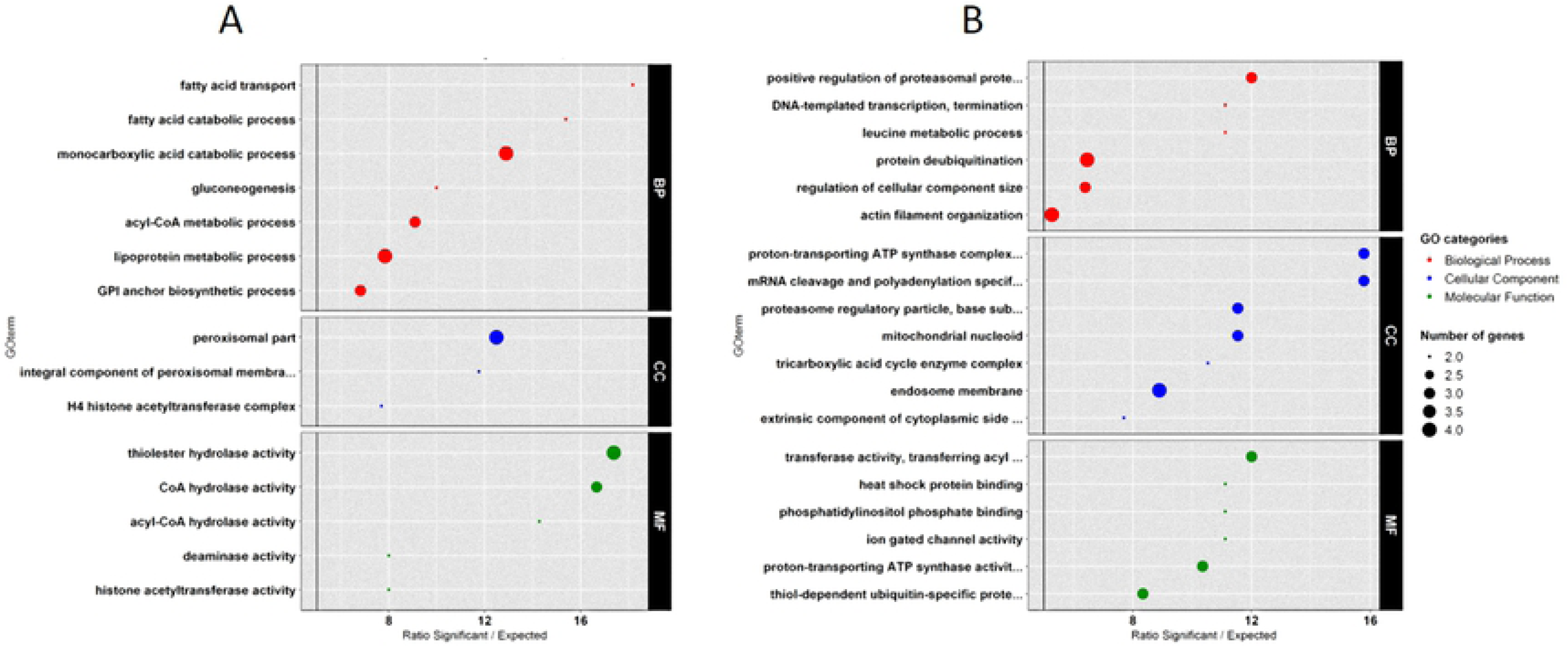
GO enrichment on DEGs explaining growth profiles. Enrichment factor (> 5) is represented on the x-axis and enriched GO-terms are described on the y-axis (A) 250 DEGs explaining PG38 growth at neutral pH (B) 351 DEGs explaining PG6 growth at acidic pH BP = Biological Process, CC = Cellular Component and MF = Molecular Function

In the same way, GO-enrichment on the 351 genes overexpressed in PG6 at acidic pH compared to neutral (**blue long arrow on Fig. 5)**, showed that 53 participated to enrich 19 GO-terms. Among these GO-terms, we particularly identified two main enriched pathways. The first one involved proteasome complex (GO-term ‘proteasome regulatory particle’) and ubiquitin carboxyl-terminal hydrolases genes (UCH): GO-terms ‘protein deubiquitination’ and ‘thiol-dependent ubiquitin-specific protease activity’. The ubiquitin-proteasome system plays a role in response to stress or protein degradation in fungi [31], and deubiquitinating enzymes (which could remove protein ubiquitination) are known to be involved in an important regulatory strategy in ubiquitin mediated protein turnover [32]. The higher growth rate at acidic pH for PG6 could be linked to a high potential of protein turnover. The second enriched pathway was linked to ‘proton-transporting ATP synthase activity’ which provides energy for ATP synthesis, directly related to RNA synthesis; this might explain in part the growth especially through high transcriptomics activity and protein synthesis.

### RT-qPCR validation of RNA-Seq analyses

Eight genes, which were significantly differentially expressed at least twice among the four main comparisons, were selected to validate the RNA-Seq data by RT-qPCR assay. RT-qPCR primers are described in **S2 table**. The selected genes take part of the Pal pathway (palB, palC, palF, and pacC) or are related to pathogenesis (lac1, lac2, gmk1, and exo) [8]. This panel provided a large range of expression levels and enabled us to compare log2FoldChange values between RT-qPCR and RNA-Seq experiments. We found a moderate correlation (r^2^ = 0.51) between log2FoldChanges of both types of experiments but with a very strong significance according to Pearson correlation test (p-value < 0.01), confirming the validity of the RNA-Seq data (**Fig. 8 A**). The similarity of the data between RT-qPCR and RNA-Seq was more precisely confirmed by comparing the pacC (GGTG_01809) expression (**Fig. 8 B**). In both types of experiments, pacC displayed similar expression’s pattern: it was less expressed at acidic pH than at neutral pH, as observed in other fungi such as *A. nidulans* [33], *Sclerotinia sclerotiorum* [34], *Fusarium Oxysporum* [35], *Colletotrichum acutatum* [36], *Aspergillus oryzae* [37], or *Coniothyrium minitans* [38].

**Fig 8.**
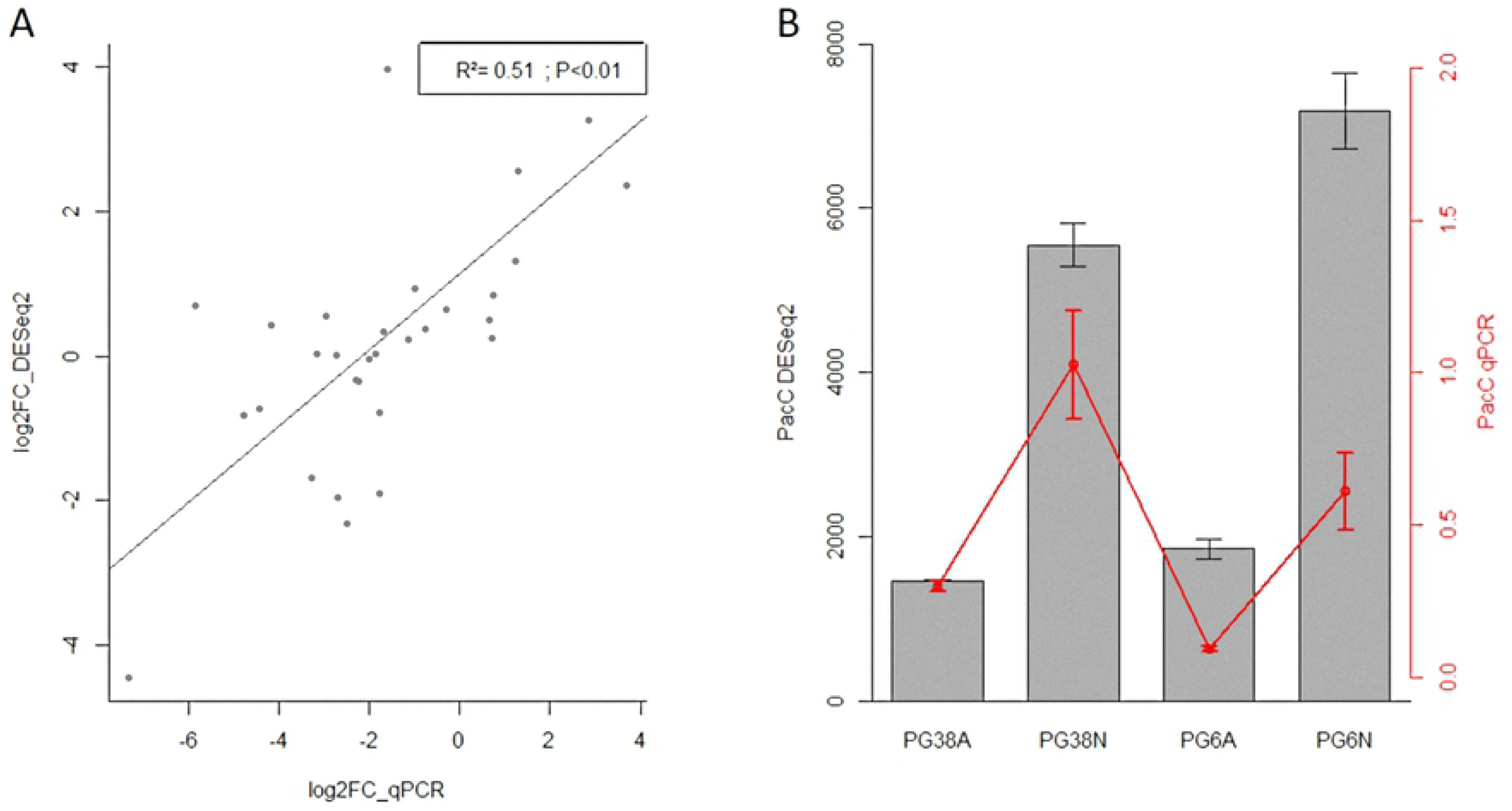
RT-qPCR and RNAseq data comparison. A. Correlation between log2FoldChange of quantitative real-time PCR (x-axis) and RNAseq (y-axis) on 8 selected genes in the four main contrasts. B. Comparison of pacC mean expression in all samples (black bars represents RNAseq results; red points represented RT-qPCR results). Errors bars represented standard errors of the means. A: acidic pH; N: neutral pH; pacC DESeq2: transcript expression of pacC (DESeq2 normalized counts); pacC qPCR: transcript expression of pacC (RT-qPCR expression normalized with external controls).

Although slight variations in correlations were observed between the two methods, similar trends in transcript abundances were generally observed confirming a reliable expression result generated by RNA sequencing.

## Conclusions

This study is gives new insights on gene expression regulation monitored by the ambient pH in a strain-specific way with opposite growth rate profiles in function of the extracellular pH.

Independently of the ambient pH, the two strains have different strategies in adapting transcriptome to growth: PG38 expressed mainly genes involved in response to stresses (DNA repair and antibiotic resistance), whereas genes involved in mitochondrial inheritance and energy transfer were more expressed in PG6. So PG38 (G2 type) could have a better ability to survive and/or to resist to stresses, whereas PG6 (G1 type) could have a better ability to grow. Moreover, both strains are able to regulate their intracellular pH and activated mechanisms to produce ascospores when growing in neutral conditions whereas they overexpressed functions linked to mitochondria transport in acidic conditions.

This study also highlighted that both strains adopted different strategies to grow, depending on the ambient pH. These two strains were able to grow on both pH conditions. At acidic pH, PG6 was able to highly grow and showed an enrichment of pathways related to protein deubiquitination and ATP synthesis, but no genes involved in survival strategies were particularly expressed. On the other hand, PG6 was disadvantaged at neutral pH because it had lower ability to grow and to survive or resist to stresses. These two characteristics could finally lead to a decrease of this type of strains in fungal population. Concerning PG38, the strain had better growth in neutral conditions, especially via the fatty acids biosynthesis pathway. At acidic pH, despite a lower growth, this strain displayed ability to stress resistance potentially linked to survivability. In total, whatever the pH but particularly at neutral pH, PG38 could have better abilities to survive in soils during intercrops, so allowing the presence of inoculum source to the next wheat culture. Thus, we described, in an original way, different pH-dependent strategies related to growth rate profiles in the two studied strains.

To go further in the understanding of survival and infection dependent on the pH and on the strain, particularly concerning the peak of G2 type after few years of wheat monoculture in neutral conditions, other similar studies including more strains and pH conditions are necessary. Moreover, *Ggt* transcriptomics study in host plant grown in soils of different pH would be interesting to focus more precisely on the infection stage. At this time indeed, only one transcriptomics study highlighted that over 3,000 genes (including genes involved in signal transduction pathways, development, plant cell wall degradation, and response to plant defense compounds) were differentially expressed between *Ggt* in culture and *Ggt* infecting roots, but this study is based on a single strain not characterized for G1/G2 type [39].

## Author contributions

| **Contributor Role** | **Role Definition** |
| --- | --- |
| **Conceptualization** | Stéphanie DAVAL |
| **Data Curation** | Joëlle AMSELEM, Kévin GAZENGEL, Nicolas LAPALU |
| **Formal Analysis** | Kévin GAZENGEL, Lionel LEBRETON |
| **Funding Acquisition** | Stéphanie DAVAL, Denis TAGU |
| **Investigation** | Stéphanie DAVAL, Kévin GAZENGEL, Anne-Yvonne GUILLERM-ERCKELBOUDT, Lionel LEBRETON |
| **Methodology** | Kévin GAZENGEL, Anne-Yvonne GUILLERM-ERCKELBOUDT, Lionel LEBRETON |
| **Project Administration** | Stéphanie DAVAL |
| **Resources** | Joëlle AMSELEM, Stéphanie DAVAL, Kévin GAZENGEL, Anne-Yvonne GUILLERM-ERCKELBOUDT, Nicolas LAPALU, Lionel LEBRETON |
| **Software** | Joëlle AMSELEM, Kévin GAZENGEL, Nicolas LAPALU |
| **Supervision** | Stéphanie DAVAL |
| **Validation** | Joëlle AMSELEM, Stéphanie DAVAL, Nicolas LAPALU |
| **Visualization** | Stéphanie DAVAL, Kévin GAZENGEL |
| **Writing – Original Draft Preparation** | Stéphanie DAVAL, Kévin GAZENGEL |
| **Writing – Review & Editing** | Joëlle AMSELEM, Stéphanie DAVAL, Kévin GAZENGEL, Nicolas LAPALU, Lionel LEBRETON, Denis TAGU |

## Supporting information captions

**S1 Table. Composition of the media used in this study**.

**S2 Table. Oligonucleotide primers used in this study**.

